# Beyond biodiversity loss defaunation: cascades effects in zoonotic disease ecology

**DOI:** 10.64898/2026.07.21.739897

**Authors:** Felipe Aramburo Jaramillo, Juan Manuel Cordovez, Carolina Bello, Mauricio Santos Vega

**Affiliations:** Grupo de Biología y Matemática Computacional (BIOMAC), Departamento de Ingeniería Biomédica, Universidad de los Andes, Bogotá D.C., COL; Departamento Ciencias Biológicas, Universidad de Los Andes, Bogotá D.C., COL; Department of Environmental System Science, Eidgenössische Technische Hochschule Zürich, Zürich, CH; Center of Ecosystem Biology, University of Rwanda, Kigali, RW

## Abstract

Defaunation, defined as the loss or severe decline of animal populations, is increasingly recognized as a major dimension of global environmental change, yet its role in zoonotic disease ecology remains poorly understood. Although defaunation significantly influences ecological dynamics, current metrics often focus on species loss without fully addressing its cascading impacts, which can affect ecological processes such as zoonotic disease transmission. In this perspective, we propose a defaunation-cascade framework linking anthropogenic pressure to zoonotic pathogen dynamics through four ecological steps: host trait filtering, abundance restructuring, interaction rewiring, and altered pathogen circulation or proliferation, and potential spillover hazard. We illustrate this framework by using a global review of empirical studies and combining georeferenced data on disease prevalence with environmental and human-related factors. Our exploratory models suggest pathogen-specific associations: parasitic and bacterial prevalence were more strongly associated with anthropogenic pressure and mammal diversity metrics, whereas viral prevalence showed weaker support. We contend that incorporating defaunation cascades into One Health surveillance will improve ecological understanding of pathogen circulation and enhance the ecological accuracy and predictive power of zoonotic disease models, thereby identifying landscapes where faunal simplification may elevate spillover hazard.

## The Emergence of defaunation and Its Functional Consequences

Over the last 500 years, human activity has led to a rapid and widespread decline in animal populations, a phenomenon that may rival past extinction events in scale and speed (Barnosky et al., 2011; Ceballos et al., 2015). This ongoing loss of animal life, increasingly referred to as defaunation, involves the decline or extinction of individuals or entire populations of animal species from natural ecosystems (Chapin et al., 2013). Defaunation is now recognized as a significant form of anthropogenic environmental change that produces biodiversity loss; however, it is not simply a reduction in species richness. Defaunation is a selective process that changes which species remain in ecosystems, their abundance, the functional composition of the community, and the ecological functions they perform (Dirzo et al., 2014; Finn et al., 2023).

Species loss is non-random and disproportionately affects large-bodied species, apex consumers, and taxa with extensive home ranges or low reproductive rates, which are often disproportionately vulnerable to hunting, habitat loss, fragmentation, and other anthropogenic pressures (Chichorro et al., 2019; Fritz & Purvis, 2010). These selective losses lead to shifts in community structure that increasingly favor smaller, fast-reproducing species, including rodents and mesopredators (Brose et al., 2017; Lee & Jetz, 2011). Defaunation therefore alters not only the number of species present, but also the distribution of functional traits, trophic roles, and life-history strategies within communities.

These changes have been documented to affect ecosystem functioning (Bogoni et al., 2020; da Silva Batista et al., 2025; Donoso et al., 2020; Galetti & Dirzo, 2013). For example, the removal of key functional groups such as large frugivores and seed dispersers can disrupt plant recruitment, forest regeneration, and carbon storage (Bello et al., 2015; Fricke et al., 2022; Pérez-Méndez et al., 2016). In parallel, the decline of apex consumers can increase herbivore and mesopredator abundances, generating trophic cascades that influence habitat structure and functional composition (Galetti et al., 2017; Young et al., 2016). Collectively, these changes modify ecosystem organization, producing cascading effects across taxonomic groups and ecosystem compartments

Recognizing defaunation as ecological reorganization, not just species loss, is crucial for understanding its broader effects. By filtering species by traits, restructuring abundances, and altering interactions, defaunation reshapes the biological context of pathogen circulation. This reframing connects faunal loss to zoonotic disease ecology by focusing on changes in function and interactions, not just species or individual counts.

## Ecological components that regulate pathogen dynamics

Despite increasing awareness of how defaunation affects ecosystem function, its potential role in influencing pathogen transmission remains poorly studied. Zoonotic diseases are infections that circulate among non-human animals and can be transmitted to humans, often through spillover events where pathogens cross species barriers from wildlife reservoirs (Rahman et al., 2020). Zoonotic pathogens persist in nature within sylvatic transmission cycles involving wildlife host species that maintain, transmit, or amplify infections over time, and, in some cases, vectors such as arthropods mediate transmission among hosts (Tompkins et al., 2011). These systems are particularly relevant to human health because zoonotic diseases account for approximately 60% of emerging infectious diseases, and more than 70% of these originate in wildlife reservoirs (Daszak et al., 2000; Woolhouse, 2002). Classic examples include leishmaniasis, cycling through wild mammals and phlebotomine sand flies; Chagas disease, sustained by wild mammal reservoirs and triatomine bugs; and yellow fever, which circulates among non-human primates via mosquito vectors before spilling into human populations (Barrett & Higgs, 2007; Coura & Viñas, 2010; Jori et al., 2013; Valentine et al., 2019).

Ecological conditions that sustain pathogen circulation rely on multiple interacting factors. The composition of the host community influences which species can be infected, whereas reservoir competence reflects how well these species acquire, preserve, and transmit pathogens (Keesing & Ostfeld, 2021a; Ostfeld & Keesing, 2000). Host abundance influences encounter rates among infected and susceptible individuals, and in vector-borne systems, vector abundance and feeding preferences shape the probability of transmission among hosts (Simpson et al., 2012; Thongsripong et al., 2021). Habitat structure, climate, and land-use conditions can further modify transmission by affecting host movement, vector breeding sites, and contact among wildlife, domestic animals, and people (Begon, 2008; Cable et al., 2017; Kilpatrick & Randolph, 2012).

Defaunation may influence disease-relevant ecological conditions by altering species composition, host availability, reservoir competence, vector densities, pathogen reproduction, and patterns of interspecific contact (Dobson, 2004; Gervasi et al., 2015; Prospero & Cleary, 2017). These shifts may reshape spillover hazard by modifying reservoir availability, host–vector contact rates, and opportunities for interaction among wildlife, domestic animals, vectors, and humans (Gibb et al., 2020; Merrill & Johnson, 2020; Ostfeld & Keesing, 2012). For instance, rodent-borne diseases have been shown to intensify in defaunated systems across multiple regions, including Africa (Young et al., 2014), North America (Dizney & Dearing, 2016), and Central America (Suzán et al., 2009). Similar patterns have been reported in both mammalian and avian disease systems in wildlife communities undergoing defaunation (Allan et al., 2009; Markandya et al., 2008; Rocha et al., 2013; Roque et al., 2013). However, these outcomes are unlikely to be universal. The effects of defaunation on pathogen dynamics depend on pathogen traits, transmission mode, reservoir competence, vector ecology, and host community structure. Robust quantitative evidence linking defaunation to disease dynamics remains limited, partly because most studies do not jointly measure faunal loss, host functional traits, community composition, interaction networks, and pathogen prevalence (Keesing & Ostfeld, 2021b; Rahman et al., 2020). Understanding how wildlife loss alters infectious disease dynamics requires more than just cataloging vanished species. We develop a framework to clarify how defaunation reorganizes host communities, affecting traits, abundances, and interactions, which influence transmission, with the aim of anticipating how faunistic change will reshape zoonotic and wildlife pathogen circulation.

## Defaunation-Driven Pathways of Zoonotic Disease hazard

Human activities such as hunting, land conversion, and habitat fragmentation are major drivers of faunal decline and reorganization, leading to cascading ecological consequences (Cardillo et al., 2006; Dirzo et al., 2014). Building on these mechanisms, we conceptualize defaunation as a cascade linking anthropogenic pressure to pathogen circulation within wildlife communities and, potentially, to increased spillover hazard. The cascade operates through four interconnected pathways. First, defaunation filters host communities by selectively removing large-bodied, slow-reproducing, disturbance-sensitive species while favoring smaller-bodied, fast-reproducing, disturbance-tolerant taxa. Second, these trait shifts restructure host abundance and dominance, increasing the relative importance of generalist species that persist in human-modified landscapes. Third, altered dominance and functional composition rewire ecological interactions, predation, competition, host–host contacts, and host–vector encounters. Fourth, these interaction changes modify pathogen circulation by shifting reservoir availability, host competence, vector feeding opportunities, and contact rates between susceptible and infected hosts.

In Figure 1, we present a schematic representation of potential relationships among anthropogenic pressures, biodiversity loss, changes in traits and composition, and host–vector–pathogen interactions. Human activity, as measured by pressure metrics such as the Human Footprint Index, is associated with habitat modification and conditions that are expected to contribute to defaunation. In transformed environments, habitat modification may increase the availability of breeding sites for vectors and reduce the abundance of predators that regulate vector populations, thereby altering host–vector contact rates (LaDeau et al., 2015; Li et al., 2014; Simpson et al., 2011). Defaunation may further weaken ecosystem functions by removing species with important regulatory or functional roles, thereby indirectly influencing pathogen transmission. For example, the loss of frugivores and seed dispersers can hinder forest regeneration and alter vegetation structure, thereby affecting habitat suitability for vectors and reservoir hosts (Chahad-Ehlers et al., 2018; Chapin et al., 2013; Zittra et al., 2017). Simultaneously, decreasing functional and phylogenetic diversity can weaken community-level processes that control transmission. Meanwhile, streamlined food webs and broken species interactions promote conditions that benefit generalist reservoir species, thereby increasing pathogen persistence and transmission risk.

The resulting shifts in community composition, encompassing traits, abundance, diversity, and the structure of host–host and host–vector interactions, modify the conditions under which pathogens circulate, altering transmission rates and pathogen reproduction within host populations (Civitello et al., 2015). Whether these changes suppress or amplify transmission largely depends on how reservoir competence is distributed within the reorganized community (Streicker et al., 2013). Disease ecology has framed this question around two non-mutually exclusive hypotheses: the dilution effect, whereby diverse host communities reduce transmission through the buffering influence of less competent species, and the amplification effect, whereby community reorganization favors competent reservoirs and intensifies pathogen spread (Pauciullo et al., 2024; Wood et al., 2014). We contend that the central question is not whether biodiversity loss increases or decreases disease risk, but how defaunation reorganizes ecological communities. The loss of particular functional groups, the persistence or expansion of others, and the resulting rewiring of species interactions can produce either dilution or amplification effects, depending on the functional identity of the species lost and those that remain. Consequently, predicting disease outcomes requires moving beyond species richness alone and examining changes in community functional structure and ecological interaction (Keesing & Ostfeld, 2021b; Ostfeld & Keesing, 2012).

For instance, in Lyme disease systems of the eastern United States, diverse vertebrate communities dilute *Borrelia burgdorferi* transmission by diverting tick feeding toward less competent hosts; yet when mammalian diversity declines, white-footed mice (*Peromyscus leucopus*) come to dominate, tipping the same system toward amplification (Keesing et al., 2010). Conversely, studies of West Nile virus transmission in avian communities in Atlanta, Georgia, found that dilution or amplification outcomes were determined more by the identity of the species present in the host community than by overall species richness, with communities dominated by less competent avian hosts showing reduced transmission despite substantial viral circulation (Levine et al., 2017). Together, these cases illustrate that the direction of the effect is not an intrinsic property of biodiversity loss itself, but a consequence of which functional groups, amplifying or diluting, are lost and which persist or expand.

The decline of top predators can trigger mesopredator release and rodent population expansion, increasing the relative abundance of competent reservoir hosts and thereby promoting pathogen amplification in disturbed systems (Young et al., 2016). These defaunation-driven shifts in host abundance, community dominance, and transmission pathways can be further intensified — or in some cases counteracted — by human-driven landscape modification, depending on how it reshapes the resulting community. In addition, agricultural expansion, urbanization, and forest fragmentation increase spatial overlap among wildlife, humans, and domestic animals (Johnson et al., 2013; Pievani, 2014). This increased interface elevates contact rates among wildlife hosts, domestic animals, vectors, and human populations, often, though not universally, in landscapes dominated by synanthropic rodents and vectors, thereby amplifying spillover potential.

Figure 1 contrasts these pathways across landscapes differing in human pressure. In less-transformed landscapes, wildlife communities retain high taxonomic diversity and large-bodied vertebrates, maintaining the ecological complexity that constrains pathogen transmission, consistent with dilution-effect dynamics. In highly transformed landscapes, defaunation restructures communities toward smaller-bodied, disturbance-tolerant species that often serve as effective reservoirs, amplifying transmission within sylvatic cycles and increasing overlap among wildlife, vectors, and people. This community reorganization also reshapes host-vector interactions: the loss of large-bodied vertebrates that typically act as poor or dead-end hosts for many vectors concentrates vector feeding on the remaining competent reservoir species, potentially increasing vectorial capacity and pathogen circulation within the community.

**Figure 1.**
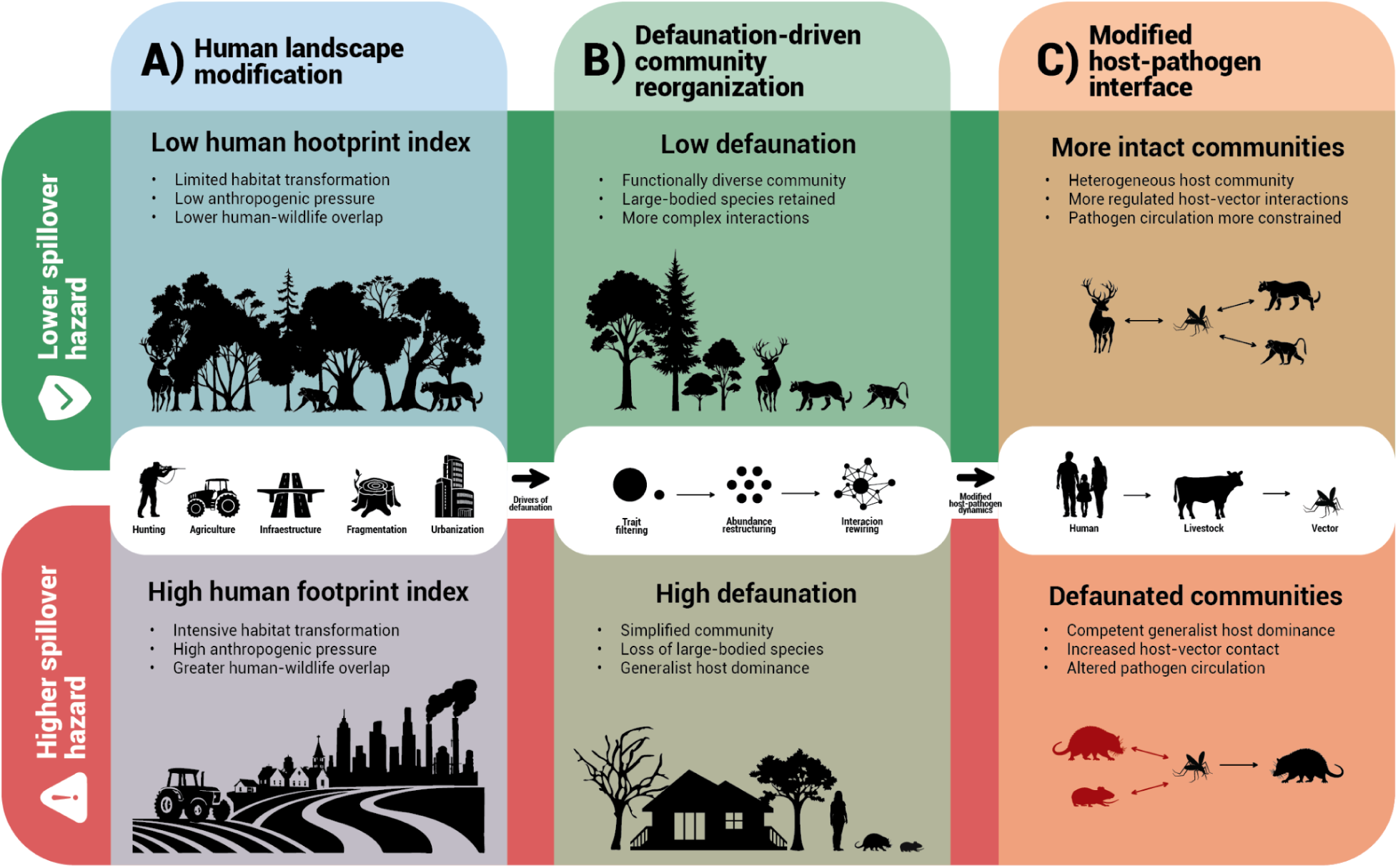
Conceptual diagram illustrating pathways linking human landscape modification, defaunation, and zoonotic disease hazard across two contrasting scenarios. (A) Human landscape modification, represented by the Human Footprint Index, generates gradients of habitat transformation and human–wildlife overlap. Low-footprint landscapes retain limited habitat transformation and lower human–wildlife overlap, whereas high-footprint landscapes are characterized by intensive transformation, high anthropogenic pressure, and greater contact between humans and wildlife. (B) These pressures drive defaunation and community reorganization through trait filtering, abundance restructuring, and interaction rewiring, shifting communities from functionally diverse assemblages that retain large-bodied species and complex interactions toward simplified systems dominated by generalist reservoir hosts. (C) Resulting changes in host composition and ecological interactions modify the host–pathogen interface: more intact communities maintain heterogeneous host composition and more tightly regulated host–vector interactions that constrain pathogen circulation, whereas defaunated communities show dominant competent hosts, increased human–wildlife contact, and altered pathogen circulation. Together, these contrasting pathways illustrate how anthropogenic disturbance may increase spillover hazard by reshaping ecological communities, host–vector dynamics, and transmission pathways.

Critically, spillover to humans does not follow from altered wildlife dynamics alone. It requires increased human contact with infected hosts or vectors, through encroachment into wildlife habitats, expansion of agricultural frontiers, or greater shared use of landscapes by people, domestic animals, and wildlife. Defaunation thus operates as an intermediary process linking anthropogenic drivers of biodiversity loss to altered host-pathogen dynamics, shaping the spillover hazard by pathogen traits and functional composition of the communities in which pathogens circulate, the spillover event will depend on the degree to which humans intersect with those communities. This framework is not a universal causal model, but it underscores the importance of incorporating the cascading effects of faunal loss into epidemiological surveillance and disease risk assessments in rapidly changing landscapes.

## Conceptual Gaps in Linking Defaunation and Zoonotic Disease Surveillance

Although the number of studies examining connections between environmental change, biodiversity loss, and human disease risk is increasing (Gibb et al., 2020; Van Uhm & Zaitch, 2021), changes in animal community composition and their interactions have rarely been explicitly incorporated into models of zoonotic disease transmission or surveillance systems. Most research continues to emphasize climate change, land-use transformation, and direct human–wildlife contact as the principal drivers of zoonotic hazards (Frumkin, 2022; Pierrehumbert, 2019), while overlooking how the removal of key wildlife groups and their functional roles can indirectly alter disease processes through ecological reorganization. This is a critical gap. Defaunation reshapes host community composition, abundance, functional traits, and interaction structure, altering transmission rates and pathogen persistence across environmental gradients (Bernardo-Cravo et al., 2020; Glidden et al., 2021). As a result, current surveillance systems may detect pathogen presence or human cases without understanding the previous ecological processes that have led to pathogen maintenance and spillover (Carlson et al., 2021; Morse et al., 2012). Recognizing defaunation as a central ecological process can therefore improve our understanding of a zoonotic hazard by highlighting the mechanisms through which biodiversity loss influences disease dynamics, including the demographic expansion of competent reservoirs, the weakening of trophic regulation, and the erosion of functional diversity that can otherwise constrain pathogen transmission (Pongsiri et al., 2009; Rohr et al., 2019).

Compounding these data gaps is a problem of interpretation. Spillover hazard is not equivalent to spillover risk (Lloyd-Smith et al., 2009; Plowright et al., 2017). Spillover hazard reflects pathogen circulation within wildlife communities, whereas spillover risk also depends on pathogen shedding, transmission route, vector ecology, livestock interfaces, human exposure, and socioeconomic context (Becker et al., 2020; Lo Iacono et al., 2016). Critically, host competence and interaction networks, the two ecological properties most directly linking community composition to transmission intensity, are rarely measured in surveillance studies (Albery et al., 2020). Spillover hazard should therefore be interpreted as one ecological component of spillover risk, not as a direct measure of risk to humans. This distinction is essential for linking defaunation to disease surveillance without overstating the implications of prevalence data.

#### Box 1. Evidence, proxies, and patterns: what a global wildlife pathogen dataset can and cannot tell us about defaunation-driven disease risk

##### Part I. Evidence, proxies, and the limits of inference

To illustrate the potential and limitations of current data and to propose mechanisms underlying defaunation in disease dynamics, we compiled studies on pathogen prevalence in wildlife and integrated them with georeferenced environmental, anthropogenic, and host-diversity data. Our aim is not to prove causality, as direct measures of faunal loss are rarely available. Instead, we assess whether wildlife pathogen prevalence varies with gradients in human activity, environment, and host diversity, using these as proxies for the defaunation-cascade framework. This offers a hypothesis-generating view, highlighting data gaps for future zoonotic surveillance.

##### Dataset structure and geographic scope

Our evidence synthesis compiled 142 georeferenced prevalence estimates from 107 empirical studies on parasitic, bacterial, and viral zoonotic pathogens in wildlife hosts (Figure 2A). Study sites cluster in North America, Europe, and parts of sub-Saharan Africa and Southeast Asia, reflecting longstanding asymmetries in surveillance capacity rather than the global distribution of disease risk. Large regions of the tropics and subtropics, where defaunation pressure tends to be highest and mammalian diversity greatest, are substantially underrepresented. This geographic skew limits the generalizability of observed associations and should be understood as a structural feature of the available evidence base, not a sampling choice.

Each prevalence estimate is a snapshot: a single host species, a single pathogen, at a single location and time. The dataset does not record local community composition, longitudinal changes in host assemblages, or direct evidence of faunal loss. It provides a set of geographic anchors that can be linked to spatially explicit ecological and anthropogenic variables, a foundation for pattern detection, not for causal attribution.

##### What the predictors actually measure

Predictor variables were grouped into three categories: environmental conditions, human disturbance, and host community structure. Environmental variables included mean annual temperature and precipitation (Brun et al., 2022), vegetation heterogeneity, aboveground biomass (Kumar & Mutanga, 2017; Tuanmu & Jetz, 2015), and biome identity, with study sites assigned to terrestrial biomes following Dinerstein et al., (2017). Anthropogenic pressure was characterized using net forest cover change (Harris et al., 2021), the Biodiversity Intactness Index (BII), and the Global Human Footprint Index (HFI; Venter et al., 2016), which integrates land use, infrastructure, human population density, and accessibility at 1-km resolution. Host community structure was characterized using potential mammalian species richness and rarity-weighted richness derived from species distribution data rather than local inventories (Albuquerque et al., 2019; Veach et al., 2017). Generalized linear mixed models (GLMMs) were used to explore associations between pathogen prevalence and these ecological gradients while accounting for variation among pathogens and biomes; full model specifications and outputs are provided in supplementary materials (Table S1).

Because direct, spatially explicit estimates of defaunation were unavailable for most study sites, and existing defaunation layers are restricted to particular regions and taxa (Benítez-López et al., 2019; Ferreiro-Arias et al., 2025), HFI was used as a proxy for anthropogenic pressures associated with defaunation. Comparisons with available regional defaunation layers confirmed positive associations between HFI values and independently derived estimates of faunal depletion. Nevertheless, HFI cannot capture hunting intensity, local extirpations, or realized changes in community composition and should be interpreted as an indirect indicator of conditions associated with defaunation rather than a direct measure of faunal loss. Similarly, mammalian species richness and rarity-weighted richness reflect the potential pool of mammals whose ranges overlap a given site, not the species actually present or detectable there. A site with high potential richness may harbour a depauperate and heavily hunted local assemblage; rarity in geographic range extent does not map onto low local abundance or high functional uniqueness in any consistent way.

##### Why does this constrain causal inference?

Taken together, these constraints mean that the analyses test whether pathogen prevalence varies along gradients of *conditions associated* with defaunation, not whether defaunation itself drives prevalence. Unmeasured confounders, host identity, reservoir competence, transmission route, sampling effort, and detection probability, are likely to explain substantial variance within and across pathogen groups. Statements of the form “defaunation elevates prevalence” or “community homogenization amplifies transmission” cannot be sustained by these data alone. The models should be read as hypothesis-generating rather than hypothesis-confirming, and the associations they identify as entry points for targeted investigation rather than evidence of specific ecological mechanisms.

Considering these constraints, namely, a geographically biased evidence base, proxy variables representing unmeasured defaunation, and cross-sectional snapshots that cannot determine causality, we focus on what the data can effectively provide: a depiction of how pathogen prevalence varies in relation to ecological gradients linked to defaunation across the 142 study sites compiled herein. Figure 2 encapsulates this evidence and the exploratory associations it identifies.

**Figure 2.**
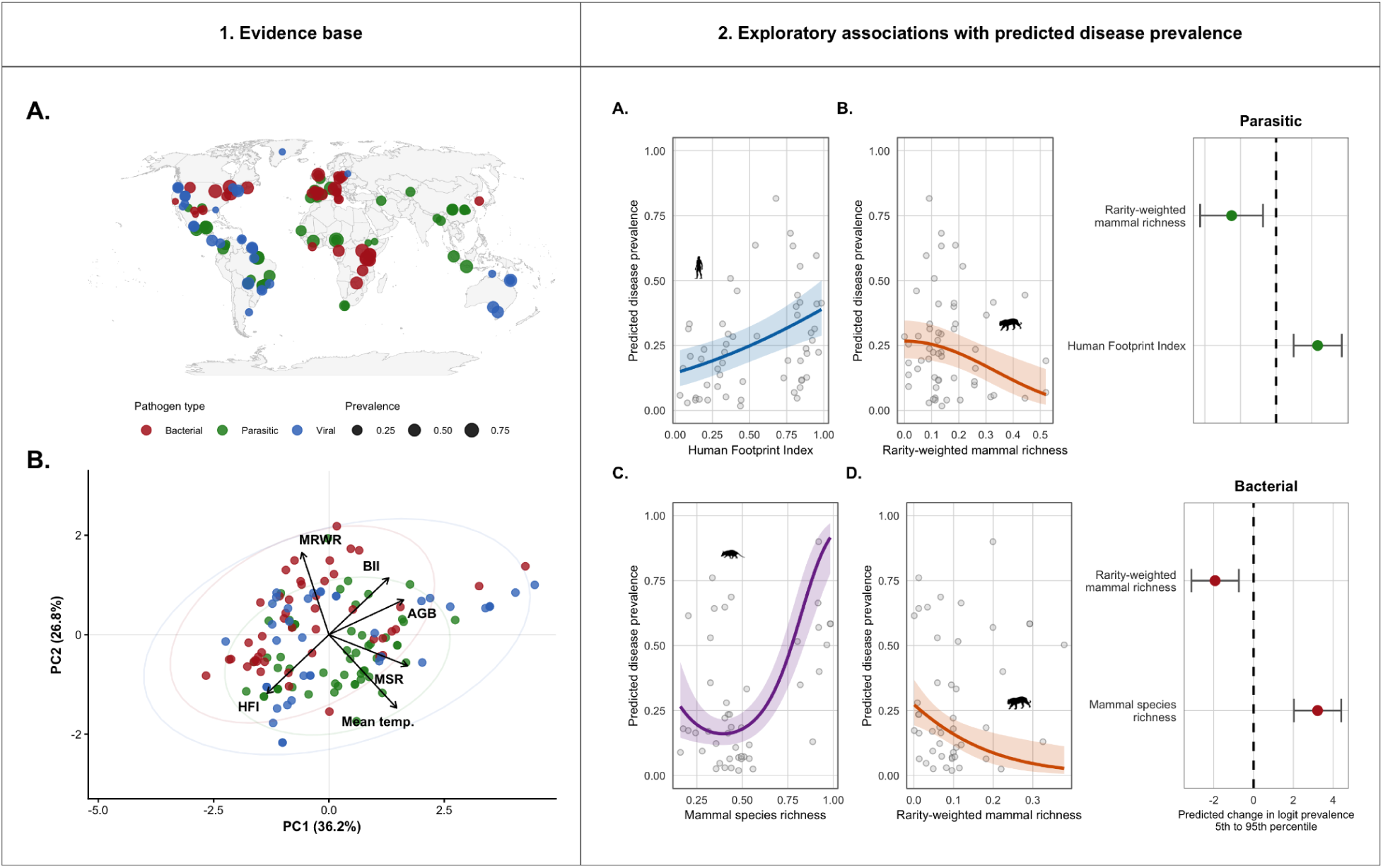
Global evidence base and exploratory associations between ecological gradients and wildlife pathogen prevalence. (A) Geographic distribution of the 142 wildlife pathogen prevalence estimates included in the evidence synthesis. Points are colored by pathogen group (parasitic, bacterial, and viral), and point size is proportional to the reported prevalence at each study location. (B) Principal component analysis (PCA) summarizing the environmental, anthropogenic, and host-community gradients represented across study sites. Vectors indicate the contribution of predictor variables used in the exploratory analyses. (C–F) Exploratory generalized linear mixed model predictions illustrating the associations between pathogen prevalence and ecological gradients retained in the best-supported models for parasitic and bacterial pathogens. For parasitic systems, prevalence was positively associated with the Human Footprint Index and negatively associated with rarity-weighted mammal richness. For bacterial systems, prevalence increased with mammal species richness and decreased with rarity-weighted mammal richness. Forest plots summarize model-averaged coefficient estimates and 95% confidence intervals for predictors retained within the best-supported models. These analyses are intended to identify broad ecological patterns and generate hypotheses rather than establish causal relationships, providing the empirical motivation for the mechanistic framework developed in subsequent sections.

**Figure 3.**
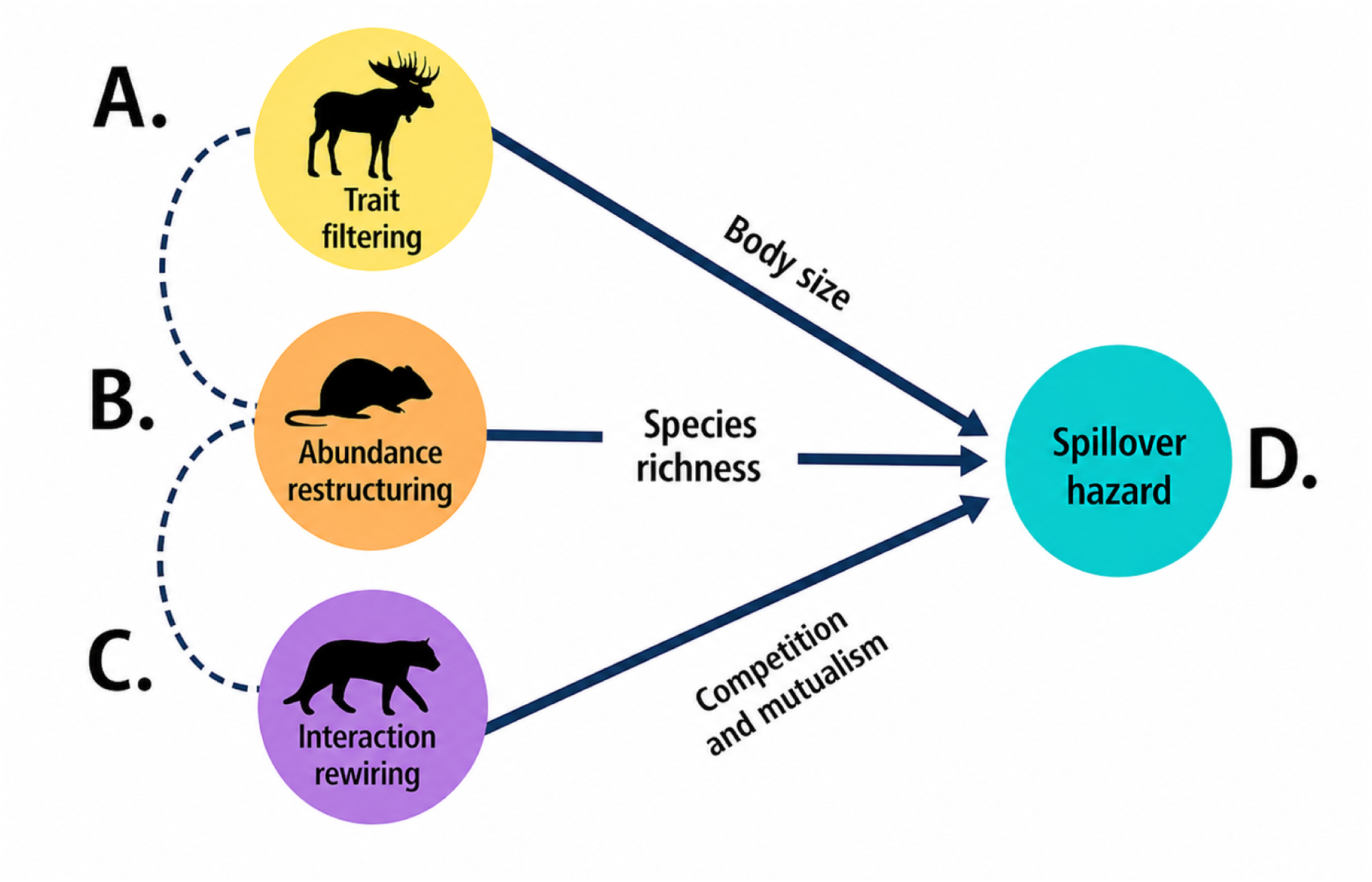
Conceptual representation of the ecological cascade through which defaunation may influence zoonotic spillover hazard. Defaunation is represented as a sequence of interconnected ecological processes rather than a single loss of species. (A) Trait filtering. Selective species loss reshapes the functional composition of wildlife communities by disproportionately affecting species with particular traits, including body size, life-history strategy, trophic position, and reservoir competence. (B) Abundance restructuring. Changes in community composition alter species dominance and relative abundance, modifying the distribution of potential reservoir hosts and overall community structure. (C) Interaction rewiring. Community reorganization changes ecological interactions among hosts, vectors, predators, and competitors, potentially altering transmission pathways and contact structure. These three processes are interdependent, with changes in functional composition influencing demographic responses, which in turn reshape ecological interactions. (D) Spillover hazard. Together, these cascading ecological processes may modify pathogen circulation within wildlife communities and alter the ecological conditions under which spillover to livestock or humans becomes more likely.

Considering these constraints, we now explore what the preliminary analyses suggest. The comprehensive dataset covers a wide geographic and ecological spectrum of wildlife pathogen studies (see Figure 2A–B). Principal component analysis reveals that study sites encompass major gradients of human impact, environmental factors, and host community composition. The first axis primarily distinguishes human pressure from biodiversity-related variables, while the second axis captures climatic differences. Overall, stronger and more consistent links were observed for parasitic and bacterial pathogens compared to viral pathogens (refer to Supplementary Tables S2 and S3). Variables related to human activity and host community structure appeared more often in the most supported models than climatic or productivity measures, underscoring ecological gradients that need deeper mechanistic research. Box 2 summarizes these findings and outlines what is necessary to advance from correlation to understanding mechanisms.

#### Box 2. Patterns, mechanisms, and what better data would require

##### Parasitic prevalence and anthropogenic pressure

Parasitic pathogen prevalence tends to increase with measures of anthropogenic pressure (HFI) and decrease with rarity-weighted mammalian richness (Figure 2, Exploratory associations, panels A–B). If this pattern reflects dilution-loss dynamics, where anthropogenic pressure removes diverse host communities and concentrates transmission on competent reservoir species, it is consistent with core predictions of dilution effect theory. The association persists after accounting for broad-scale climatic variation, suggesting it is not simply a latitudinal artifact driven by the co-distribution of human pressure and tropical species richness.

##### Bacterial prevalence and host community structure

Bacterial prevalence showed stronger associations with metrics describing the potential mammalian host community, including overall species richness and rarity-weighted richness (Figure 2, Exploratory associations, panels C–D). Sites with higher potential host diversity tend to show lower bacterial prevalence, consistent with inter-host dilution; sites where rarity-weighted richness is reduced show elevated prevalence, suggestive of homogenization-driven amplification. This cannot be confirmed, however, without local species inventories that capture realized rather than potential community composition.

##### Viral prevalence: no consistent ecological signal

Viral pathogens showed little evidence of consistent relationships with any of the ecological gradients considered. This may reflect genuine biological heterogeneity, viruses vary enormously in host range, transmission mode, and environmental persistence, or limited statistical power given the smaller number of viral observations and the absence of transmission-route covariates. The null result cautions against assuming that the ecological gradients relevant for macroparasites and bacteria generalize uniformly across pathogen types.

Taken together, these findings indicate that the relationship between habitat change, host diversity, and pathogen prevalence is neither linear nor universal, but system-specific and mediated by the functional composition of the host community, underscoring the importance of moving beyond species counts toward metrics that capture who is lost, not only how many.

###### What better data would allow us to test

Three categories of data would substantially increase the inferential leverage of future syntheses.

##### Local community inventories paired with prevalence estimates

Studies that measure realized host assemblages, species identities, relative abundances, and functional traits — at the same sites and times as pathogen prevalence would allow direct tests of diversity–prevalence relationships and distinguish dilution from amplification at the community level. This would replace the current reliance on range-map potential (Albuquerque et al., 2019; Veach et al., 2017) with ecologically grounded measures of who is actually present and at what density.

##### Reservoir competence and host functional traits

Linking host species to empirical estimates of competence and trait databases covering body mass, life-history pace, immune parameters, and habitat specialization would transform species richness counts into functional metrics directly relevant to transmission dynamics. It would allow the question to shift from how many species are present? to what is the functional composition of the host community, and how does it change as defaunation proceeds?

##### Longitudinal surveillance at defaunation fronts

Before–after or along-gradient designs at sites with documented hunting pressure, habitat conversion, or wildlife extraction would provide the temporal contrast needed to move from correlation to causal inference. Cross-sectional prevalence snapshots, however numerous, cannot establish whether changes in community composition preceded or followed changes in pathogen prevalence. Prospective monitoring at active defaunation fronts, particularly in regions currently under-represented in global surveillance datasets, is the most direct path to mechanistic understanding.

Until such data are available, the patterns identified here serve as an empirical scaffold: consistent enough to motivate targeted surveillance and to anchor the conceptual framework developed below, but not sufficient to adjudicate among competing mechanistic explanations. The processes we describe, dilution loss, community homogenization, and competence amplification, remain open hypotheses whose confirmation requires the kinds of data that wildlife disease ecology has rarely prioritized collecting.

These broad-scale patterns serve as an empirical point of departure for the conceptual framework developed below, which focuses on how changes in community composition, species abundances, and ecological interactions may influence pathogen circulation in defaunating landscapes. Rather than testing specific hypotheses about transmission processes, the analysis identifies ecological pathways that warrant further investigation and highlights key data gaps that limit our understanding of how defaunation shapes zoonotic disease dynamics.

## Cascading Mechanisms: Evidence and Knowledge Gaps

The patterns described above suggest that anthropogenic pressure and host community structure are associated with pathogen prevalence in system-specific ways, but they do not reveal the processes responsible for those associations. Both the studies included in our synthesis and our own analysis rely largely on broad-scale proxies of environmental change and community structure, while direct measurements of host traits, species abundances, reservoir competence, and interaction networks remain scarce. Consequently, the mechanisms most likely to link wildlife community change to pathogen dynamics are often inferred rather than measured, limiting our ability to move from pattern to process.

A first mechanism involves host trait filtering. Defaunation often occurs unevenly across species, with large-bodied species being disproportionately lost from communities (Cardillo et al., 2006; Chichorro et al., 2019; Tomiya, 2013). Body size is a crucial trait in this context because it correlates with life-history strategies, population density, and immune investment, all of which influence host competence and transmission potential (Downs et al., 2019; Silk & Hodgson, 2021). Smaller-bodied species generally have faster life histories, higher population densities, and greater tolerance to infection, traits linked to increased pathogen richness and reservoir competence (Albery & Becker, 2021). Yet most disease-prevalence datasets do not jointly measure host traits, local faunal loss, and reservoir competence, limiting our ability to test whether trait filtering is actually responsible for observed changes in pathogen prevalence.

A second mechanism involves host abundance and dominance. As a result, the selective loss of large-bodied species shifts community composition toward smaller, generalist taxa, many of which serve as effective pathogen reservoirs (Plourde et al., 2017; Young et al., 2016), and this change is often accompanied by increased abundance and dominance of rodents/bats and other opportunistic species in disturbed landscapes (Keesing & Ostfeld, 2024). These shifts can change community evenness and the distribution of reservoir competence, thereby influencing whether biodiversity loss is associated with dilution or amplification. However, abundance data are rarely available in global wildlife disease datasets. As a result, analyses often rely on species richness or potential distribution layers, which cannot distinguish between a species-rich assemblage with low reservoir dominance and a simplified assemblage dominated by a few competent hosts.

A third mechanism involves interaction rewiring. Defaunation affects interactions between hosts and vectors by decreasing trophic regulation and competition, thus increasing encounter rates between them. This change can facilitate pathogen transmission and spillover in specific ecological contexts (Judson et al., 2016; Plowright et al., 2024). These network-level changes may strongly influence pathogen circulation, but they are rarely measured in surveillance studies. Without data on host–host contacts, vector feeding preferences, predator regulation, and wildlife–livestock–human interfaces, it remains difficult to determine whether defaunation alters pathogen prevalence through interaction pathways rather than through richness or abundance alone.

These limitations are also geographic. The empirical research is geographically biased, with most studies on defaunation and disease dynamics focusing on North America and Europe, while tropical regions that are biodiversity hotspots and that concentrate the higher burden of zoonotic disease (Allen et al., 2017), remain underrepresented. This geographic imbalance is also apparent in our dataset and limits the general applicability of the current conclusions. This imbalance is problematic because tropical landscapes often combine high host diversity, high pathogen diversity, rapid land-use change, hunting pressure, and limited disease surveillance capacity. Improving geographic representation would reduce sampling bias and improve the generality of model-based inferences about defaunation and pathogen circulation.

These levels are not independent: trait distributions constrain demographic responses, demographic shifts alter interaction strengths, and interaction networks feed back on population sizes and trait selection. Disease prevalence emerges from the coupling of all three. The distinction between solid and dashed arrows reflects an important caveat: while the ecological mechanisms are conceptually well-grounded, their connection to defaunation is often inferred from coarse proxies, introducing uncertainty into causal attribution. Zoonotic risk under defaunation should therefore be understood as the outcome of interacting changes across the trait, demographic, and network dimensions of community structure, rather than the consequence of any single ecological variable.

## Ecological data priorities for zoonotic surveillance

To move beyond broad-scale associations and better understand how community change influences pathogen circulation, future surveillance efforts should prioritize ecological data that are both difficult to obtain and directly relevant to transmission processes. While environmental variables such as climate, vegetation structure, land cover, and human pressure can increasingly be retrieved from global spatial datasets, information on wildlife infection, host communities, and ecological interactions remains scarce. Particularly important are geo-referenced estimates of pathogen prevalence in wildlife hosts, accompanied by information on host identity, host abundance or relative detection rates, local community composition, and vector occurrence where applicable. Data describing host–vector interactions, including blood-meal analyses and feeding preferences, would provide critical insight into transmission pathways that are currently inferred rather than measured. Likewise, information on host competence, or functional proxies associated with competence, would help clarify how community composition translates into pathogen circulation. Improved characterization of wildlife–livestock–human interfaces is also needed to understand how changes in animal communities influence spillover hazard.

Emerging technologies offer new opportunities to collect these data at broader spatial scales. Environmental and invertebrate-derived DNA (eDNA/iDNA), metabarcoding approaches, camera traps, acoustic monitoring, blood-meal analyses, and other molecular tools can help characterize wildlife communities, species interactions, and pathogen occurrence without requiring prohibitively intensive field campaigns. Combined with remotely sensed environmental information, these approaches may help bridge the gap between broad-scale ecological patterns and the biological processes that underpin pathogen transmission.

Ultimately, progress in understanding the links between defaunation and zoonotic disease will depend less on developing additional environmental covariates than on improving our ability to measure wildlife infection, community composition, species interactions, and transmission-relevant ecological traits. Expanding these data across currently underrepresented tropical regions will be particularly important for building more mechanistic and ecologically grounded frameworks for zoonotic surveillance in rapidly changing landscapes.

## Measuring community reorganization across the defaunation cascade

Although environmental datasets describing climate, land use, vegetation structure, and anthropogenic pressure have become increasingly available at global scales, the ecological information required to understand how wildlife community change influences pathogen transmission remains remarkably limited. Consequently, most studies—including our own exploratory analysis—relate pathogen prevalence to broad environmental gradients without directly measuring the biological processes through which defaunation reshapes disease dynamics. The principal limitation is therefore no longer the availability of environmental predictors, but rather the scarcity of ecological data describing how animal communities are reorganized following biodiversity loss.

Understanding these processes requires moving beyond static descriptions of environmental change toward direct measurements of community reorganization. Defaunation is not simply the disappearance of species; it is a progressive restructuring of ecological communities that alters functional composition, demographic structure, species interactions, and ultimately pathogen circulation. Capturing this cascade therefore requires increasingly detailed ecological information together with analytical frameworks capable of representing processes operating across multiple organizational levels. Rather than advocating a single modeling framework, we propose a progression from increasingly informative ecological data to increasingly mechanistic inference (Table 1).

**Table 1.** Roadmap for progressing from broad ecological patterns to mechanistic understanding of defaunation and zoonotic disease dynamics. Each level builds upon the previous one by combining increasingly informative ecological data with analytical frameworks that support progressively stronger ecological inference.

| Level | Defaunation pathway | Ecological process | Key ecological data | Analytical framework | Mechanistic insight |
| --- | --- | --- | --- | --- | --- |
| A. | Trait filtering | Selective loss or persistence of species with particular functional traits (body size, trophic position, life history, reservoir competence). | Species occurrences, functional trait databases, local community composition. | Trait-based models; functional diversity analyses. | Reveals how selective species loss restructures the functional composition of animal communities. |
| B. | Abundance restructuring | Changes in abundance, dominance and demographic turnover following selective species loss. | Camera traps, occupancy surveys, abundance estimates, eDNA/iDNA, metabarcoding, passive acoustic monitoring. | Hierarchical community models; abundance-based transmission models. | Identifies how demographic restructuring changes reservoir availability and transmission potential. |
| C. | Interaction rewiring | Reorganization of host–host, host–vector, predator–prey and competitive interactions. | Blood-meal analyses, telemetry, movement ecology, co-occurrence data, ecological interaction networks. | Multilayer network models; movement models; dynamic transmission models. | Explains how community reorganization reshapes transmission pathways through ecological networks. |
| D. | Spillover hazard | Changes in pathogen circulation across wildlife–livestock–human interfaces. | Geo-referenced wildlife prevalence, vector occurrence, livestock distribution, human exposure and contact data. | Spatial spillover models; integrated surveillance; early-warning frameworks. | Identifies where community reorganization is most likely to increase zoonotic spillover risk. |

The first level of inference concerns trait filtering. Defaunation selectively removes species according to characteristics such as body size, trophic position, life-history strategy, and reservoir competence (Cardillo et al., 2006; Han et al., 2016; Ripple et al., 2014). Integrating species occurrence records with functional trait databases allows researchers to quantify how community composition changes under anthropogenic pressure. Trait-based models and functional diversity analyses provide the first opportunity to move beyond species counts and identify which ecological functions are disappearing from communities (Gibb et al., 2020; McGill et al., 2006).

The second level focuses on abundance restructuring. Defaunation rarely removes species uniformly; instead, it alters their relative abundance and dominance, often favouring disturbance-tolerant generalists while reducing large-bodied or specialised taxa (Dirzo et al., 2014; Young et al., 2016). Camera traps, occupancy models, eDNA/iDNA, metabarcoding and passive acoustic monitoring increasingly allow these demographic responses to be quantified and incorporated into hierarchical community and abundance-based transmission models.

The third level addresses interaction rewiring. Changes in species composition and abundance reorganize host–host, host–vector and predator–prey interactions, yet these processes remain among the least frequently measured components of wildlife disease ecology. Blood-meal analyses, movement ecology, telemetry, co-occurrence analyses and ecological interaction networks provide complementary information on transmission pathways and support multilayer network and dynamic transmission models (Borremans et al., 2019; Delmas et al., 2019; Oliveira-Santos et al., 2021; Sah et al., 2017).

The final level links these ecological processes to spillover hazard. Integrating wildlife prevalence with vector occurrence, livestock distribution, human exposure and wildlife–livestock–human interfaces enables spatial spillover models and early-warning systems capable of identifying landscapes where anthropogenic pressure, community reorganization and opportunities for cross-species transmission converge. Progress therefore depends less on increasingly sophisticated statistical models than on collecting the ecological data needed to parameterize and validate mechanistic frameworks across diverse ecological contexts.

## Integrating cascading effects into One Health approach

The One Health framework recognizes that human, animal, and environmental health are fundamentally interconnected (Adisasmito et al., 2022; Mackenzie & Jeggo, 2019). Yet, in practice, One Health surveillance often emphasizes human cases, livestock outbreaks, pathogen detection, and broad environmental drivers such as land-use change or climate. Although these components are essential, they rarely capture the ecological reorganization of wildlife communities that precedes pathogen emergence. Incorporating the defaunation cascade into One Health therefore represents an opportunity to shift attention from the consequences of spillover to the ecological processes that shape zoonotic risk before spillover occurs.

A defaunation-aware One Health approach expands surveillance beyond pathogen detection by incorporating information on wildlife community structure, including functional composition, species abundances, ecological interactions, and wildlife–livestock–human interfaces. Rather than treating biodiversity simply as a background environmental variable, this framework recognizes community reorganization itself as an important component of disease risk. Integrating these ecological measurements with pathogen prevalence, vector occurrence, and human exposure data would improve our ability to identify landscapes where anthropogenic pressure is restructuring wildlife communities in ways that may favor pathogen circulation.

The framework proposed here also highlights an important distinction between environmental drivers and ecological responses. Climatic variables, land-use change, and remotely sensed indicators of habitat structure are increasingly available through global datasets (Brun et al., 2022; Kumar & Mutanga, 2017), but they cannot directly quantify local faunal loss, changes in species dominance, or the reorganization of ecological interactions. These biological responses ultimately determine how pathogens circulate within wildlife communities and therefore represent the critical missing component of many current surveillance systems.

Advances in wildlife monitoring now make this transition increasingly feasible. Camera traps, environmental and invertebrate-derived DNA (eDNA/iDNA), metabarcoding, acoustic monitoring, blood-meal analyses, and movement data provide new opportunities to measure community composition, abundance, and ecological interactions at spatial scales previously unattainable. Coupled with ecological network analyses, hierarchical community models, and spatially explicit surveillance, these approaches offer the possibility of moving from broad environmental correlations toward mechanistic understanding of how community reorganization influences pathogen dynamics.

Ultimately, the contribution of the defaunation-cascade framework is not to replace existing One Health approaches but to strengthen them by explicitly incorporating ecological mechanisms linking biodiversity change to zoonotic disease dynamics. By integrating progressively richer ecological information into surveillance and modeling, One Health can move from predominantly reactive detection of outbreaks toward a predictive framework capable of identifying where community reorganization is likely to increase opportunities for pathogen circulation and spillover. Recognizing defaunation as a process of ecological restructuring, rather than simply species loss, therefore provides both a conceptual advance and a practical roadmap for anticipating zoonotic emergence while promoting the conservation of ecological integrity.

## Data availability

The data and code supporting the findings of this study are publicly available at: https://github.com/felipearamburojaramillo/Beyond-biodiversity-loss-defaunation-cascades-effects-in-zoonotic-disease-ecology.git

