## Supplementary material for "Beyond biodiversity loss defaunation: cascades effects in zoonotic disease ecology": Suplementary material

### Literature search strategy and inclusion criteria

To assemble a global dataset linking defaunation to zoonotic disease dynamics, we conducted a structured literature search designed to capture empirical studies reporting pathogen prevalence in wildlife host populations. The scope of the search was restricted to studies that quantified infection prevalence in non-human animals across parasitic, bacterial, and viral systems and provided sufficient spatial information to integrate with environmental and anthropogenic datasets.

The literature search was conducted using the Web of Science Core Collection, complemented by targeted searches in Google Scholar and Elsevier (ScienceDirect), following established guidelines for structured evidence assessments (Burton et al., 2007; Varker et al., 2015). In Web of Science, search strings were constructed by combining terms related to zoonotic or wildlife disease, biodiversity loss and defaunation, and infection prevalence using Boolean operators. These included variations of "zoonosis", "defaunation" or "biodiversity loss", and "prevalence" or "infection rate", applied across titles, abstracts, and keywords. Searches were not restricted by publication year but were limited to peer-reviewed articles in English. Additional studies were identified through backward and forward citation tracking of relevant publications.

Initial inclusion criteria required that studies be based on empirical data, employ an experimental or quasi-experimental design comparing treatments with varying degrees of defaunation, report host disease prevalence across treatments, and provide precise, georeferenced data. Because few studies explicitly incorporated defaunation as a treatment, we conducted an additional targeted search focusing on host disease prevalence across the three pathogen groups and subsequently assessed whether each study site was subject to defaunation pressure using independent spatial proxies (see main Methods). Studies reporting only human disease prevalence were excluded to maintain a focus on ecological processes within wildlife communities. Review articles, theoretical studies, and studies lacking quantitative prevalence estimates were also excluded.

All retained studies were screened in two stages (title and abstract screening followed by full-text review), and relevant data were extracted in a standardized format, including pathogen type, host species, sample size, number of infected individuals, and geographic coordinates. This process ensured consistency across studies and enabled the construction of a spatially explicit dataset suitable for comparative and modeling analyses.

**Supplementary Table S1.** Summary of environmental, structural, human-impact, and host community predictors included in the generalized linear mixed models used to evaluate the influence of defaunation on zoonotic pathogen prevalence across biomes. Colors correspond to the conceptual components in Figure 1, representing environmental context (gray), ecosystem structure (green), human pressure (blue), and host community composition (yellow), which together capture the cascading pathways linking anthropogenic change to pathogen dynamics.

| Variable | Units | Source | Category | Description |
|---|---|---|---|---|
| Mean annual rain | mm | CHELSA-bioclim (Brun et al., 2022) | Environmental | Mean annual rainfall (mm) recorded in the year the study was conducted. |
| Mean annual temperature | °C | CHELSA-bioclim (Brun et al., 2022) | Environmental | Mean annual temperature (°C) recorded in the year the study was conducted. |
| Above-ground biomass | Gt | CEDA (Kumar & Mutanga, 2017) | Structural | Above-ground plant biomass (Gt) estimated for the study year. |
| Forest cover | Mha | Global Forest Watch (Townshend et al., 2012) | Structural | Total area covered by forests, expressed in million hectares (Mha), measured for the year of the study. |
| Human footprint index | % | SEDAC (Scholes & Biggs, 2005) | Human impact | Composite index quantifying the extent of human influence on natural ecosystems, combining factors such as built environments, population density, infrastructure, and land use intensity. |

| | | | | |
|---|---|---|---|---|
| Biodiversity intactness index | % | SEDAC (Venter et al., 2016) | Human impact | Index estimating how much of a region's original species abundance persists despite human impacts, expressed as a percentage of the pre-impact state. |
| Mammal species richness | NA | IUCN<br><br>(Ceballos & Ehrlich, 2006) | Host composition | Count of distinct mammal species present within the study area or sampling unit during the year of observation. |
| Mammal weighted mammal species richness | NA | IUCN (Ceballos & Ehrlich, 2006) | Host composition | Mammal weighted mammal species richness is a measure of mammal diversity that gives greater weight to species with smaller geographic ranges. It is calculated by summing the inverse of each species' range size across all species present in a grid cell, so areas containing more range-restricted species receive higher values. |

**Exploratory mixed-effects analyses of ecological gradients associated with wildlife pathogen prevalence**

To evaluate associations between defaunation-related processes and variation in zoonotic pathogen prevalence, we fitted generalized linear mixed models (GLMMs) following Zuur et al. (2009). Pathogen prevalence was defined as the proportion of infected individuals in sampled wildlife populations and modeled with a binomial error distribution and a logit link function. Because direct measures of realized local faunal loss were rarely available across studies, this analysis uses available proxies to characterize defaunation pressure, treating the results as a basis for identifying the ecological gradients along which prevalence varies and for grounding the hypotheses developed in subsequent sections.

A global modeling approach was adopted to evaluate patterns across disease systems while accounting for ecological and pathogen-specific heterogeneity. To account for differences among ecological regions and pathogens, disease identity was included as a random effect nested within biome, allowing baseline prevalence to vary among diseases within each biome. This random-effects structure captures variation in pathogen biology, transmission mode, and host specificity while reducing non-independence across studies. Predictor variables were grouped into three conceptually distinct categories: anthropogenic pressure associated with defaunation, host community context, and environmental conditions.

Anthropogenic pressure associated with defaunation was represented by the Human Footprint Index (HFI) and the Biodiversity Intactness Index (BII), which reflect broad gradients of human disturbance. Host community context was characterized using mammal species richness (MSR) and rarity-weighted mammal richness (MRWR), derived from IUCN expert range maps. These metrics describe the potential mammalian species pool at each site rather than realized local assemblages and were therefore not interpreted as direct measures of defaunation. Environmental predictors included above-ground biomass (AGB) and mean annual temperature. In formal terms, the number of infected individuals $y_{i\square\square}$ was modeled as a binomial random variable:

$$y_{i\square\square} \sim \text{Binomial}(n_{i\square\square}, \mu_{i\square\square})$$

where $n_{i\square\square}$ is the number of individuals sampled for observation k of disease j in biome i, and $\mu_{i\square\square}$ represents the expected prevalence. The expected prevalence was connected to the predictor variables using a logit link function:

$$\text{logit}(\mu_{i\square\square}) = \beta_0 + \sum_{\square} \beta_{\square} X_{\square,i\square\square} + b_{i\square}$$

where $\beta_0$ is the intercept, $\beta_{\square}$ are fixed-effect coefficients for predictor m, $X_{\square,i\square\square}$ represents the value of predictor m for observation k of disease j in biome i, and $b_{i\square}$ is the random intercept for disease j nested within biome i. All continuous predictors were modeled using second-order polynomial terms to accommodate nonlinear relationships with pathogen prevalence.

Model selection was performed using the dredge function in the MuMIn R package, ranking candidate models by Akaike Information Criterion (AIC) and retaining those with ΔAIC < 5 (Anderson & Burnham, 2004; Harrison et al., 2018). Given the number of predictors and potential collinearity, model averaging was used to identify predictors consistently associated with pathogen prevalence across systems (Barton & Barton, 2015). In addition to the global analysis, separate models were fitted for viral, bacterial, and parasitic pathogens, using the same predictor structure and biome as a random-effects specification, to assess whether associations varied across major pathogen groups.

The final dataset comprised 142 observations from the structured review, spanning 56 parasitic, 46 bacterial, and 40 viral disease systems. The dataset encompasses zoonotic pathogens, broadly defined, including both direct transmission and vector-borne systems, provided that prevalence was estimated in wildlife host populations. The predominance of parasitic infections reflects uneven study effort across pathogen types rather than epidemiological prevalence alone.

Model selection followed an information-theoretic approach based on AIC, with inference drawn from the relative support of competing models rather than null-hypothesis significance testing. Models for parasitic and bacterial systems retained meaningful ecological and environmental predictors within the best-supported set (ΔAIC < 2; Table 1. SM), suggesting that variation in prevalence within these groups is structured along gradients of anthropogenic pressure, potential host community composition, and environmental conditions. For viral diseases and the global dataset, best-supported models were null or included only weakly supported predictors, indicating that the proxies available in this dataset do not adequately capture the drivers of viral prevalence, likely reflecting the greater dependence of viral transmission on fine-scale host contact rates, vector ecology, and immune dynamics that are rarely measured at the scales considered here.

These results guided the mechanistic hypotheses developed in subsequent analyses. The model-supported associations between ecological gradients and prevalence in parasitic and bacterial systems provide the empirical basis for the pathways illustrated in Figure 2, where anthropogenic pressure and host community restructuring emerge as the variables most consistently linked to variation in wildlife pathogen prevalence. Viral systems, by contrast, point to the need for finer-scale data on interaction networks and transmission ecology before analogous patterns can be detected.

These patterns prove to be pathogen-specific. For parasitic diseases (Figure 2, Evidence base, panels A–B), prevalence increases with human footprint but declines with higher rarity-weighted mammal richness. This is consistent with anthropogenic pressure favoring parasite circulation in simplified, homogenized communities, whereas compositionally distinctive assemblages, in which rare and functionally unique species remain present, are associated with lower prevalence. Together, these relationships suggest that both the intensity of human disturbance and the erosion of host community distinctiveness jointly shape parasite transmission, supporting the hypothesis that defaunation-driven homogenization creates conditions favorable to parasite persistence.

For bacterial diseases (Figure 2, Evidence base, panels C–D), the pattern diverges: prevalence increases sharply with overall mammal species richness but again decreases with rarity-weighted richness. This indicates that bacterial circulation benefits from a broad pool of available hosts, yet is dampened in communities where rare species contribute meaningfully to composition. In more homogeneous assemblages dominated by widespread generalist hosts, transmission appears to intensify, suggesting a role for community evenness and functional identity in regulating bacterial dynamics beyond simple species counts.

The forest plots (Figure 2, Evidence base, Parasitic and Bacterial) synthesize model estimates across systems, reinforcing that anthropogenic pressure and host community structure are the variables most consistently associated with variation in prevalence. Taken together, these findings indicate that the relationship between habitat change, host diversity, and pathogen prevalence is neither linear nor universal, but system-specific and mediated by the functional composition of the host community. This underscores the mechanistic relevance of defaunation as a driver of a zoonotic hazard and the importance of moving beyond species counts toward metrics that capture who is lost, not only how many.

**Supplementary Table S2.** Beta generalized linear mixed model examining parasitic pathogen prevalence as a function of Human Footprint Index (HFI) and mammal rarity-weighted richness (MRWR).

| **Predictor** | **Estimate** | **SE** | **95% CI** | ***p*** |
|---|---|---|---|---|
| Intercept | -1.680 | 0.280 | (-2.229, -1.130) | <0.001 |
| Human footprint index | 1.347 | 0.365 | (0.633, 2.061) | <0.001 |
| Mammal rarity-weighted richness | -6.353 | 2.029 | (-10.329, -2.377) | 0.002 |
| **Random effects** (Intercept SD) | | 0.447 | | |
| Observations | | 56 | | |
| Disease:Biome (Groups) | | 11 | | |
| Dispersion parameter | | 8.35 | | |
| AIC | | -57.3 | | |
| BIC | | -47.1 | | |
| Log-likelihood | | 33.6 | | |

*Estimates are reported on the logit scale of the beta regression. The model included a random intercept for the interaction between disease type and biome (Disease:BIOME). Confidence intervals were calculated as Wald 95% confidence intervals.*

**Supplementary Table S3.** Beta generalized linear mixed model examining bacterial pathogen prevalence as a function of mean annual temperature, mammal rarity-weighted richness (MRWR) and mammal species richness (MSR).

| **Predictor** | **Estimate** | **SE** | **95% CI** | ***p*** |
|---|---|---|---|---|
| Intercept | -1.636 | 0.207 | (-2.042, -1.229) | <0.001 |
| Mean annual temperature | -0.374 | 0.170 | (-0.707, -0.041) | 0.027 |
| Mammal rarity-weighted richness | -0.618 | 0.214 | (-10.329, -2.377) | 0.004 |
| Mammal species richness | 0.623 | 0.207 | | 0.0003 |
| **Random effects** (Intercept SD) | | 0.0005 | | |
| Observations | | 46 | | |
| Disease:Biome (Groups) | | 8 | | |
| Dispersion parameter | | 5.98 | | |
| AIC | | -41.615 | | |
| BIC | | -28.814 | | |
| Log-likelihood | | 27.8 | | |

*Estimates are reported on the logit scale of the beta regression. The model included a random intercept for the interaction between disease type and biome (Disease:BIOME). Confidence intervals were calculated as Wald 95% confidence intervals.*

**Residual-Based Assessment of Human Pressure Effects Beyond Host Diversity**

To evaluate whether human pressure explains variation in pathogen prevalence beyond that accounted for by the set of predictors retained in the final model, we implemented a residual-based analytical approach grounded in the best-supported bacterial beta regression. Specifically, we first fitted a global model including environmental, diversity, and anthropogenic predictors, and selected the most parsimonious model using an information-theoretic framework (AIC-based model selection). Prevalence was modeled using a beta distribution with a logit link, and values were transformed using a standard adjustment to ensure they lay strictly within the open interval (0,1).

Response residuals were then extracted from the best-supported model and interpreted as the unexplained component of variation in prevalence after accounting for the combined effects of the selected predictors. These residuals were subsequently modeled as a function of human pressure to assess whether any additional structure associated with anthropogenic disturbance remained. To allow for potential nonlinear relationships, we compared alternative functional forms, including linear, quadratic, and spline regressions, and selected the best-supported model using Akaike's Information Criterion (AIC).

This analysis was conducted using two complementary metrics of human pressure: the Human Footprint Index (HFI) and the Human Modification Index (HMI) (Figure S1). For HFI, the best-supported residual model was quadratic, suggesting a weak, non-monotonic pattern with a slight increase in residual prevalence at intermediate levels of human pressure. For HMI, the best-supported model was linear, indicating a marginal negative trend. However, in both cases, the fitted relationships were shallow and associated with wide confidence intervals, and residuals remained centered around zero across the full range of values.

This approach differs from traditional partial regression analyses in that it evaluates the contribution of human pressure only after accounting for the full multivariate structure identified in the best model, rather than isolating individual predictors a priori. As such, it provides a conservative test of whether anthropogenic pressure explains residual variation in prevalence beyond that already captured by ecological and environmental covariates. The weak and inconsistent patterns observed across both indices suggest limited evidence for an independent effect of human pressure on bacterial prevalence. Instead, these results are consistent with the interpretation that anthropogenic effects are primarily expressed indirectly through their influence on host community structure, reinforcing the role of defaunation and compositional change as key drivers of pathogen dynamics.

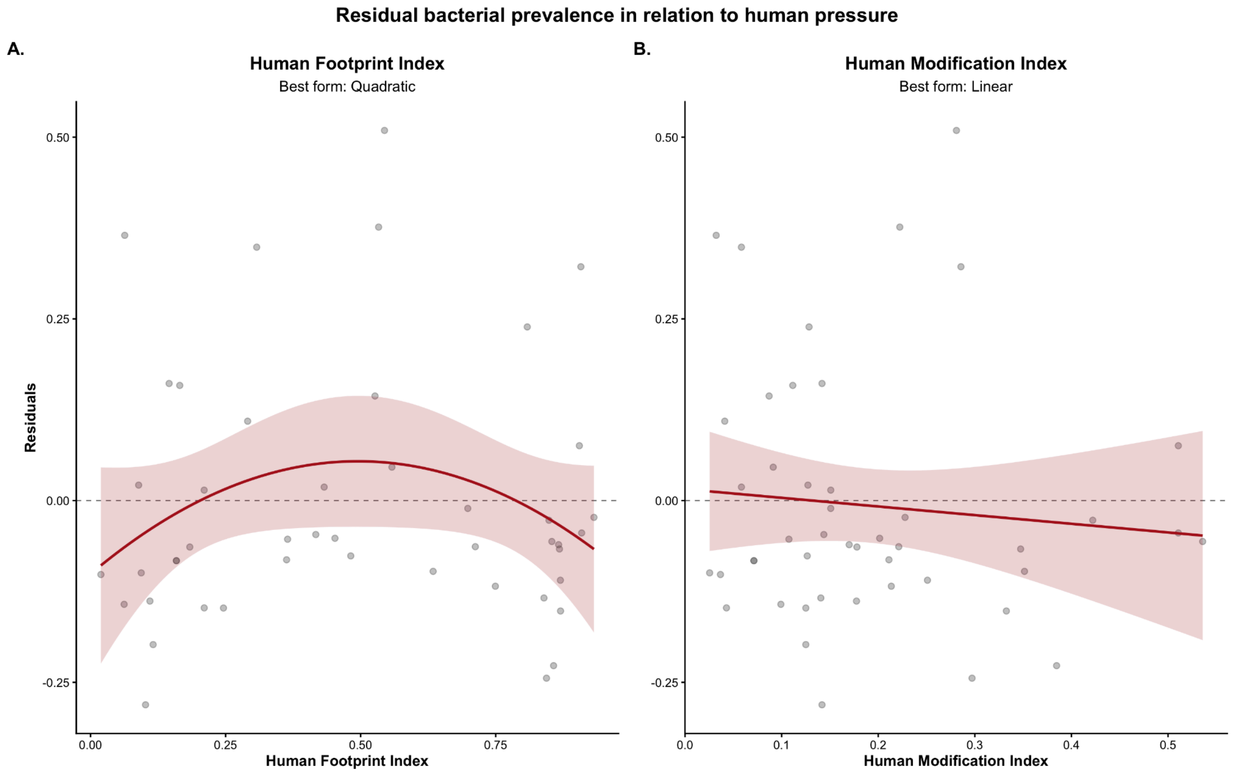


**Figure S1. Residual bacterial disease prevalence in relation to human pressure.** Residual prevalence values were extracted from the best-supported bacterial beta regression models, which accounted for the ecological and environmental predictors retained through model selection. Panel A shows the relationship between response residuals and the Human Footprint Index (HFI), for which the best-supported residual model was quadratic. Panel B shows the corresponding relationship with the Human Modification Index (HMI), for which the best-supported residual model was linear. Points represent response residuals, red lines indicate the fitted relationships, and shaded areas denote 95% confidence intervals. The dashed horizontal line marks zero residual prevalence, corresponding to the expected value under the fitted model. In both cases, the weak residual structure suggests limited evidence that human pressure explains additional variation in bacterial prevalence beyond that already captured by the selected predictors, consistent with the possibility that anthropogenic effects are primarily expressed indirectly through changes in host community structure.

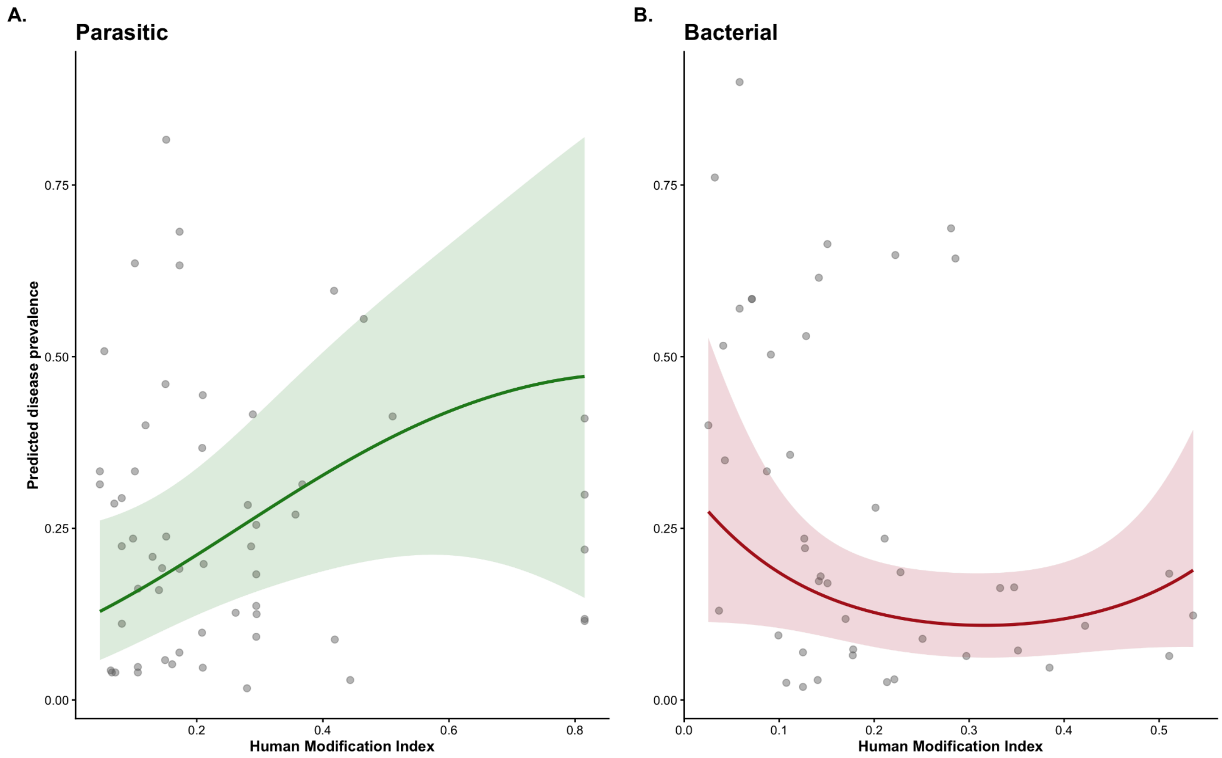


**Figure S2. Quadratic relationship between human modification and disease prevalence for parasitic and bacterial pathogens.** Points show observed prevalence values, solid lines show fitted values from separate beta regression models including human modification and its quadratic term, and shaded areas indicate 95% confidence intervals. Models were fitted separately for parasitic and bacterial datasets with disease nested within biome as a random effect.

To assess the relationship between anthropogenic pressure and disease prevalence independently of diversity-based residual approaches, we fitted beta regression models directly relating prevalence to the Human Modification Index (HMI) for each pathogen group. Models included both linear and quadratic terms for HM to allow for non-linear responses, and were fitted separately for parasitic and bacterial datasets. To account for hierarchical structure in the data, disease identity was nested within biome as a random effect. Predicted prevalence values were generated across the observed range of HM while holding other variables constant, and are presented with 95% confidence intervals. This analysis mirrors the structure of previous models but does not rely on residualization, allowing for a direct evaluation of how human-driven environmental modification shapes disease prevalence. Although the fitted relationships suggest a weak positive trend for parasitic diseases and a non-linear pattern for bacterial diseases, these effects were not statistically significant, indicating limited evidence for a consistent association between human modification and disease prevalence in these models.

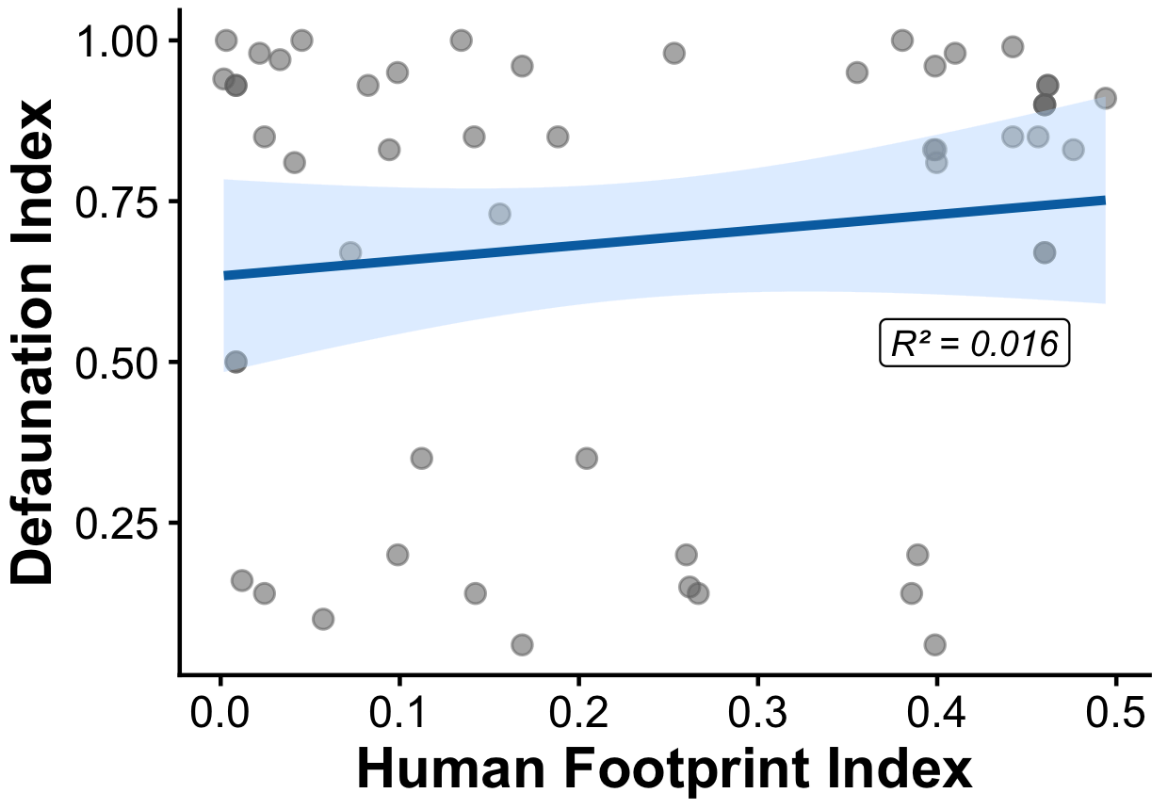


**Figure S3. Relationship between Human Footprint Index and Defaunation Index.**
Linear regression showing the association between Human Footprint Index (HFI) and the Defaunation Index across study sites. HFI is used as a proxy for anthropogenic pressure because it integrates multiple dimensions of human landscape transformation, including infrastructure, land conversion, population density, and accessibility, all of which can contribute to wildlife decline and local species loss. Under this framework, higher human footprint values are expected to be associated with greater defaunation through habitat degradation, altered species interactions, and reduced persistence of disturbance-sensitive taxa. Although the observed relationship was weak in this dataset ($R^2 = 0.016$), the figure is included to illustrate the conceptual overlap between anthropogenic disturbance and defaunation processes and to justify the use of HFI as an indirect indicator of faunal depletion.
